# Going around phase defects: reliable phase mapping for realistic data

**DOI:** 10.64898/2026.02.02.703232

**Authors:** Bjorn Verstraeten, Sebastiaan Lootens, Robin Van Den Abeele, Viktor Van Nieuwenhuize, Arstanbek Okenov, Sander Hendrickx, Arthur Santos Bezerra, Timur Nezlobinskii, Vineesh Kappadan, Balvinder Handa, Fu Siong Ng, Mattias Duytschaever, Nele Vandersickel

## Abstract

Phase mapping is a widespread method for identifying rotational electrical activity sustaining cardiac arrhythmias. However, conventional implementations assume that the cardiac phase map is continuous, leading to ill-defined phase indices in regions affected by functional conduction block, fibrosis, or anatomical boundaries. These regions of discontinuous or undefined phase, termed phase defects, lead to both false positive and false negative detections of rotational drivers. This work introduces an improved phase mapping implementation termed *extended phase mapping* that explicitly detects and accounts for phase defects, enabling robust calculation of the phase index around them. Extended phase mapping is applied to (1) simulated excitation patterns using the Fenton–Karma model, (2) experimental optical mapping data of rat ventricular fibrillation, and (3) a clinical CARTO activation map of atrial tachycardia. Across all datasets, the extended approach eliminates erroneous detections and resolves previously missed rotations. Our results demonstrate that proper treatment of phase defects yields a unified and physiologically consistent characterization of all rotational drivers including near-complete and anatomical reentries. Therefore, we propose replacing the classical notion of phase singularities with critical phase defects as the fundamental entities governing rotational dynamics in cardiac tissue.

**Author summary:** Detecting rotating electrical activity in the heart is crucial for understanding and treating abnormal heart rhythms. A common method, phase mapping, assigns a timing phase to each region of the heart to identify these rotations. However, in regions affected by scars, blocked conduction, or anatomical boundaries, the phase can become undefined or discontinuous. These so-called phase defects make current methods unreliable, causing false detections or missed rotations. In this study, we introduce an extended method that explicitly identifies phase defects and calculates phase indices around them. We test this approach using computer simulations, experimental recordings from animal hearts, and clinical heart-mapping data. Across all datasets, it eliminates false detections and reveals previously overlooked rotational activity. By properly accounting for phase defects, the extended phase mapping method provides a more reliable and complete characterization of heart rhythms, offering a physiologically meaningful framework for studying electrical dynamics in cardiac tissue.

## Introduction

Cardiac arrhythmias are a major contributor to global mortality. They remain difficult to treat effectively due to patient-specific mechanisms, with rotational electrical activity identified as a key driver [1–3]. Detecting and characterizing such activity has therefore been a central goal in cardiac simulations, experimental, and clinical electrophysiology.

In order to detect rotational activity in optical maps of a rabbit heart, phase mapping was introduced [4]. This method converts the cardiac signal to a phase, and finds infinitesimal loops where the phase differences add up to 2*π*. Since then, numerous papers have been published that use phase mapping for analyzing different datasets ranging from simulated [5–8] to experimental [9, 10] and clinical [11–16] data. These studies have demonstrated both the promise of phase mapping and its limitations when confronted with realistic data. The identification of phase singularities using phase mapping can be distorted by several physiological factors, such as functional conduction blocks [7, 8, 16, 17], scar tissue [7, 8] and anatomical structures [16]. In particular, functional conduction blocks have been recognized as equally important as phase singularities [18, 19] in describing cardiac arrhythmia dynamics. Likewise, anatomical structures play a critical role, especially in the context of ablation therapy of atrial tachycardia [20]. Together, these features underscore the need to reconsider how phase mapping algorithms handle such factors.

Our central claim is that phase mapping is conceptually sound, but naive implementations fail when confronted with discontinuous or unexcitable regions in the phase map. To demonstrate this, two algorithmic approaches are compared: a naive approach that assumes the phase map is continuous except for a finite number of singular points, and an extended approach that explicitly accounts for phase defects [21], defined as regions of discontinuous or undefined phase resulting from collisions with refractory tissue, i.e. functional conduction block, or anatomical and structural obstacles that prevent conduction in certain directions.

Both approaches are applied to three simulated activation patterns to isolate different arising problems, to an experimental optical mapping recording of ventricular fibrillation in a rat heart, and to a clinical activation map of atrial tachycardia to demonstrate the efficacy in a realistic setting. Combined, these results show that improper handling of phase defects can lead to misleading interpretations, while careful treatment restores the reliability of phase mapping.

## Materials and methods

Numerous methods exist to detect rotational activity [22–24], making it impractical to assess the impact of phase defects on all of them. Therefore, we implemented two representative approaches to phase mapping: a *naive approach*, representative of existing methods in the literature that assume the phase map is continuous except at isolated singularities, and a novel *extended approach* developed in this work, which explicitly accounts for phase defects.

Both approaches are applied to study cases of (1) simulated activation patterns around phase defects to isolate specific problems, (2) an experimental optical mapping recording of ventricular fibrillation in a rat heart to illustrate their impact in a semi-controlled setting, and (3) a clinical CARTO activation map of atrial tachycardia to evaluate the methods in realistic clinical conditions.

### Study cases

#### Finitewave simulations

We conducted proof-of-concept simulations on an isotropic 2D grid using Finitewave (https://github.com/finitewave/Finitewave), an open-source Python package for cardiac electrophysiology simulations, based on finite-difference methods, whose transparent implementation of cardiac models facilitates verification of the simulation dynamics.

We used the Fenton-Karma cell model [25] with the MLR-I parameter set, a three-variable phenomenological model suitable for generating phase defects [26] while remaining computationally efficient. Each simulation started from a mesh of 10 cm x 10 cm, with a spatial resolution of 0.25 mm. The tissue was pre-paced with 10 line stimuli (10 cm x 1.25 mm) applied from the left border of the tissue at intervals of 170 ms to establish a stable initial state with an action potential duration of 170 ms. At 1700 ms, the state of the tissue was saved and used as the starting state for the following simulations.

To investigate distinct activation patterns, we designed three simulation scenarios with specific tissue modifications and additional stimuli, depicted in Fig 1.

**Fig 1.**
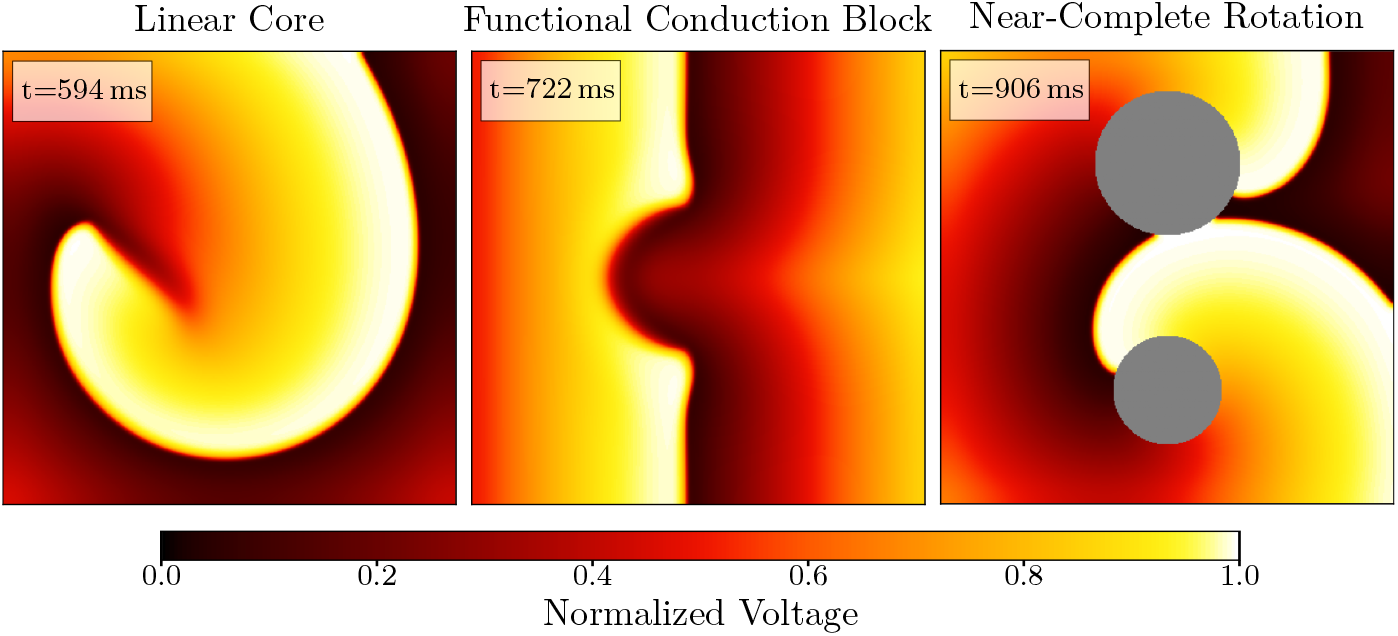
Normalized transmembrane voltage snapshots. Snapshot of the normalized transmembrane voltage in the three simulations (Linear Core, Functional Conduction Block, Near-Complete Rotation). The gray areas indicate unexcitable tissue.

- **Linear rotor core**. A single additional square stimulus was applied in the top-left corner of the tissue at 0 ms, so that this stimulus collided with the refractory tissue left by the last planar wave, producing a unidirectional conduction block. This scenario was chosen to illustrate that a linear core may appear as a meandering rotor, and the apparent complexity can be resolved once phase defects are properly accounted for.
- **Parallel activation around a functional conduction block**. Multiple line stimuli (10 cm x 1.25 mm) were applied from the left border at times: 0, 160, 315, 470, 625, 780, 935, 1090, and 1245 ms. To generate a functional conduction block in the tissue, a small central rectangular region of 5 cm by 4 mm was stimulated in between two of the line stimuli at 170 ms. The timing is such that the rectangular region is refractory when the planar stimulus reaches this area, resulting in a conduction block. This simulation represents the phenomenon of an excitation wave encountering a conduction block. This scenario contains no rotation. However, it is a common occurrence in realistic data and therefore worth investigating.
- **Complete and near-complete rotation around scar tissue**. Two circular regions of non-excitable tissue (“scar tissue”) are added to the tissue, one with a radius of 1.2 cm and one with a radius of 1.6 cm and their centers separated by 5 cm. An additional 5 cm x 5 cm stimulus was applied at 0 ms on the left side of the tissue, positioned such that it extends horizontally from the left boundary toward the center of the tissue. The stimulus is vertically centered so that it bridges the gap between both unexcitable regions. The timing of this square stimulus was chosen such that the square stimulus collides with the refractory tissue caused by the last wave, resulting in a unidirectional conduction block, located in the gap between the non-excitable regions. The other side of the square stimulus is excitable, thereby initiating a reentry pair. This scenario illustrates a regular arrhythmia characterized by a complete reentry loop around the smaller, lower obstacle and a near-complete reentry loop around the larger, upper obstacle, representing the dynamics of an atrial tachycardia (AT).

#### Optical mapping of experimental ventricular fibrillation

This paper used optical mapping data of ventricular fibrillation in a rat heart that was previously analyzed in [16] (See Fig 2). The data was obtained from [27], where details of heart preparation and ethics are described. We analyzed one case with maximally increased gap junction coupling (80 nM rotigaptide) to test our methods on stable rotors.

**Fig 2.**
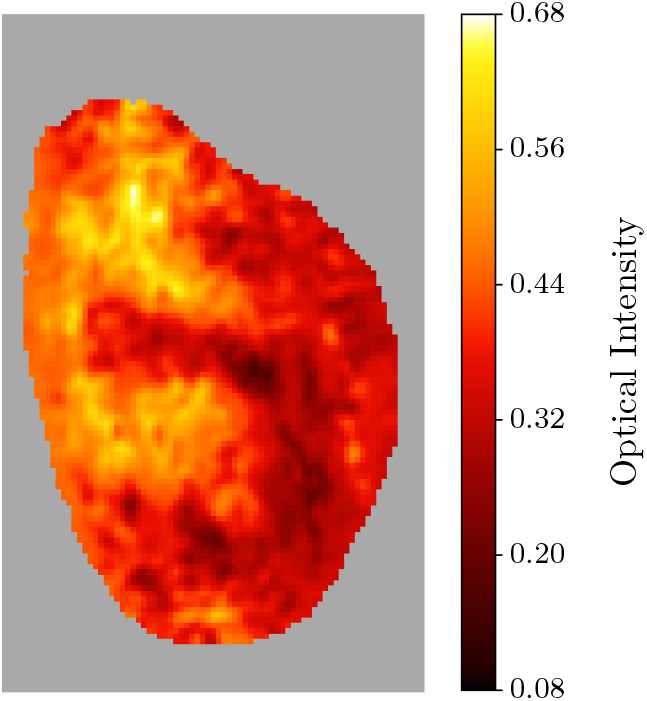
Ventricular fibrillation optical map. Snapshot (t=293 ms) of an optical intensity map of ventricular fibrillation of the left ventricle of a rat heart acquired in Imperial College London, Great Britain. The gray area indicates the background.

Briefly, a 250–300 g (9–12 weeks) Sprague–Dawley rat heart was explanted and perfused *ex vivo* via Langendorff perfusion with a Krebs–Henseleit Buffer at 37 ± 0.5^°^C and pH 7.35 ± 0.05. After 15 min, RH237 (40 µL, 5 mg/mL in DMSO) and Blebbistatin (30 µM then 10 µM) were added, plus 80 nM rotigaptide to increase gap junction coupling. VF was induced electrically at the left ventricle base (MicroPace, USA) and maintained with Pinacidil (30 µM). Optical mapping used RH237 excited at 530 nm; emission was recorded by high-resolution CMOS cameras at 1 kHz, 128 × 80 pixels, and 100–125 µm pixel width.

Data were preprocessed in MATLAB, using: background removal, 3 × 3 binning, 50-order FIR low-pass at 100 Hz, and 100 *ms* window drift removal. Further details on the signal processing protocol can be found in [28]. Afterwards, the phases were smoothed by transformation to the complex vectors 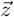 (Eq 1), and smoothing those complex vectors using a Gaussian kernel with a standard deviation of 1 pixel (100–125 µm). Finally, the resulting vectors were converted back to phases.

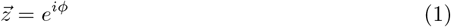

It is important to note that the measured optical intensity averages and smooths the transmembrane voltages of small regions of cardiac cells, similar to the electrogram.

#### CARTO map of clinical atrial tachycardia

The final case study is a unipolar electroanatomic map from a clinical left-atrial atrial tachycardia (AT) case recorded with the CARTO (Biosense Webster, Johnson & Johnson) system at AZ Sint-Jan Hospital, Bruges (octarray catheter), and shown in Fig 3. The map included three major anatomical boundaries: the mitral valve (MV), and the left and right pulmonary veins (LPV and RPV). This retrospective study had patient consent for scientific use. The case had the following properties:

- Whole-atrium coverage with local point density > 0.4 points/cm^2^
- Mean electrode–mesh distance < 4 mm
- Known locations of all anatomical boundaries
- Previous unsuccessful ablation of the mitral isthmus (between the MV and LPV) and the vein of Marshall
- Confirmed diagnosis of reentry around the MV + LPV and anterior scar
- Successful ablation between an anterior scar and MV, terminating the tachycardia
- Known tachycardia cycle length (TCL) before and after ablation: 317 ms to 0 ms

**Fig 3.**
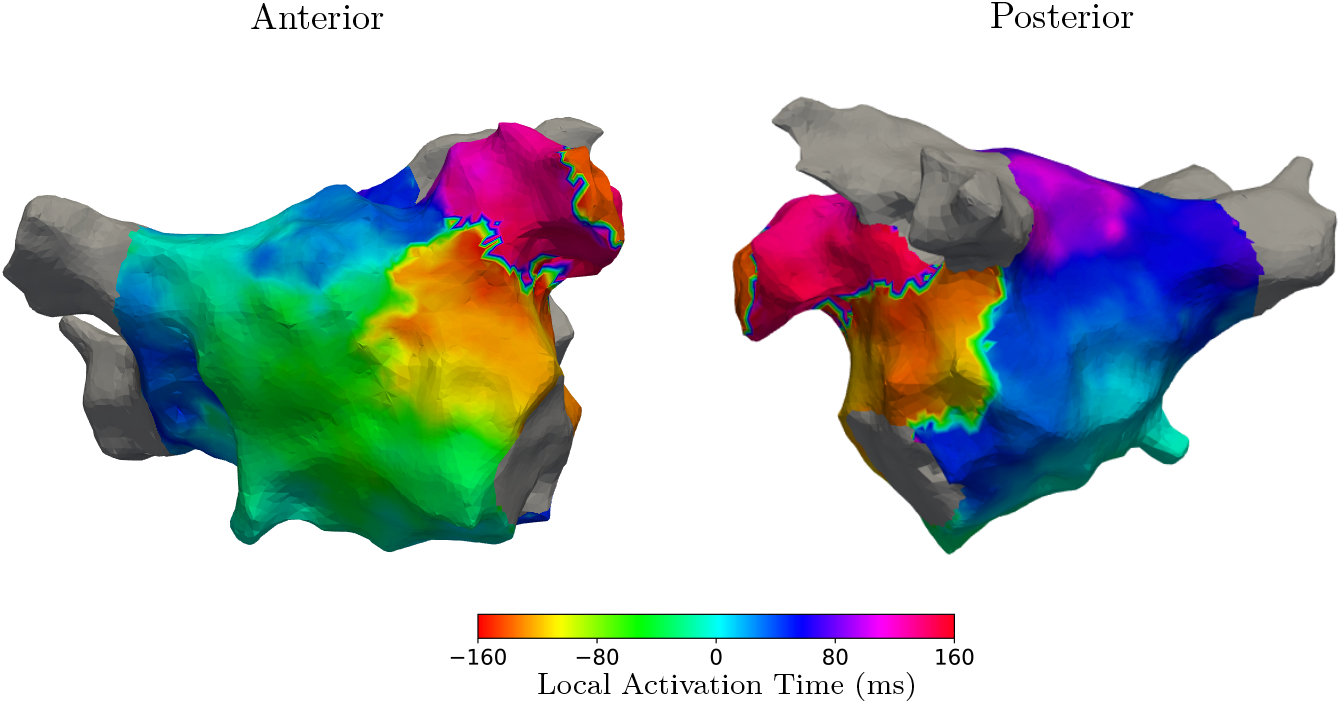
Left atrial tachycardia activation map. CARTO activation map of a clinical atrial tachycardia case of the left atrium of a patient in AZ Sint-Jan Bruges, Belgium. Both the anterior and posterior viewpoints are depicted. The gray areas indicate the anatomical valves.

The CARTO activation map data format consists of a triangular mesh of the atrium surface, consisting of points and triangles. Additionally, the scalar values on each point correspond to the interpolated local activation time (LAT) value based on the measured LAT values from the nearby electrodes. Due to the highly varying LAT values near the wavefront line, interpolation artifacts arose in the electroanatomical map. Therefore, the LAT map was preprocessed before analysis via conversion to complex vectors *z* (Eq 1) as described in the previous section, where the kernel standard deviation was 1 *mm*, and the phases are calculated via:

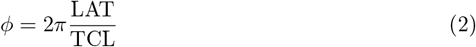

### Phase mapping

The study cases described above are analyzed using phase mapping [29], a technique commonly employed to investigate activation dynamics in cardiac tissue. The method involves constructing a phase map that represents the state of the tissue, followed by scanning the entire tissue to compute phase indexes and identify rotational activity. This process naively assumes that the phase map is continuous throughout the tissue, and is therefore referred to as the *naive approach*. In this work, a second method was developed by introducing an additional step before the computation of the phase indices. This step identifies regions where the assumption of a continuous phase field is violated. This extended procedure is referred to as the *extended approach*.

#### Phase

In cardiology, the state of a cardiac cell changes repeatedly: a cell transitions from a resting state to an excited state and eventually repolarizes back to rest. Due to this repeating behavior, the state of a cardiac cell can be approximated by a phase *ϕ*(*t*) ∈ [−*π, π*) [29].

In this paper, it is assumed that cardiac cells are activated when their measured voltage crosses a certain threshold or their time-derived voltage is maximal and crosses a threshold. Additionally, we enforce a constant rate of change in the phase between subsequent activations. As such, we construct a phase map *ϕ*(**r**, *t*), which is the collection of phases on the cardiac tissue. We calculate the phase via the sawtooth method based on LATs, illustrated in Fig 4.

**Fig 4.**
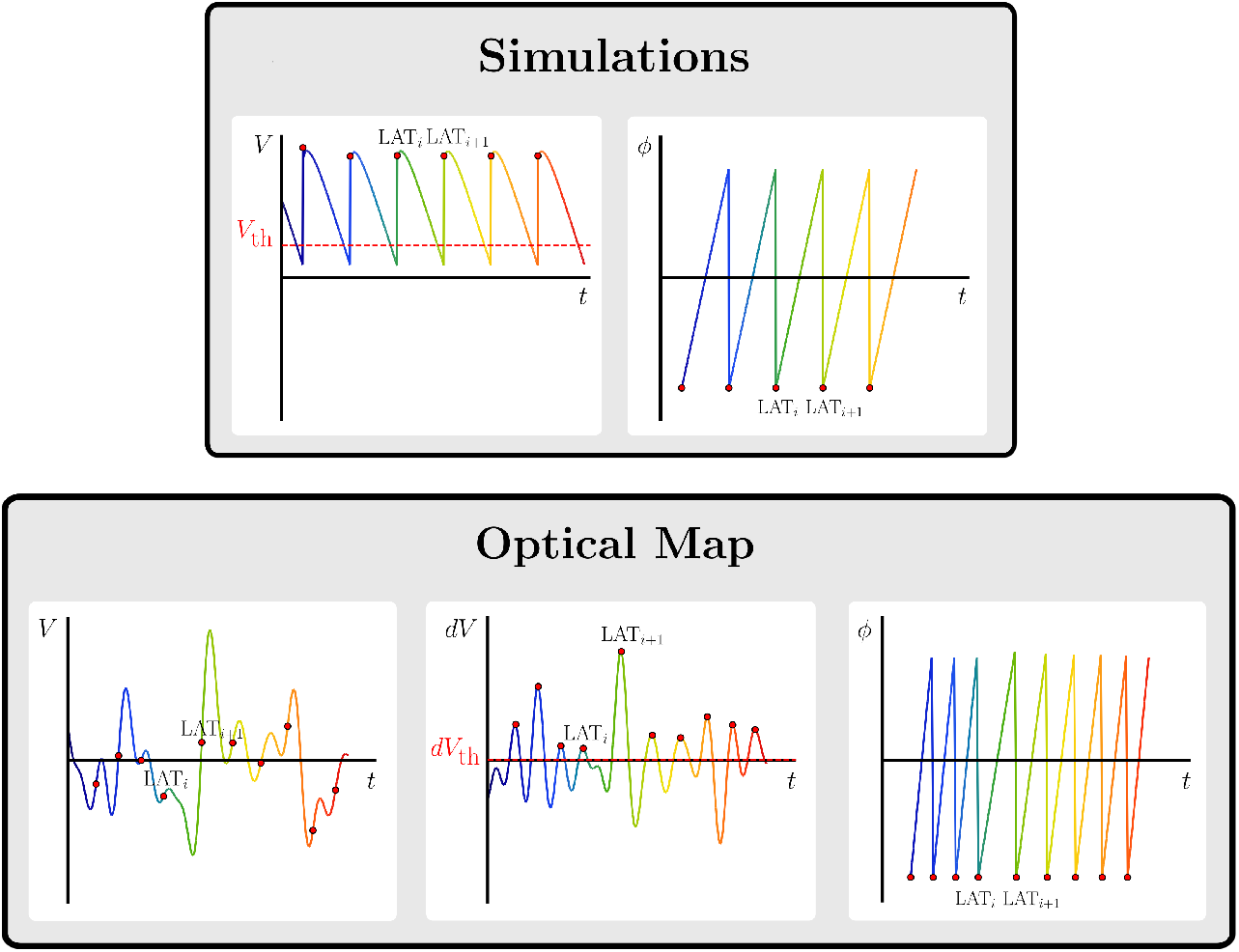
Phase computation in simulated and optical signals. *Simulations:* The first panel illustrates a simulated transmembrane voltage signal along with its annotated LATs calculated using a voltage threshold. The second panel depicts the resulting phases of the simulated voltage signal. *Optical Map:* The first panel shows an optical mapping voltage signal. The second panel illustrates the derivative of the optical mapping signal *dV* along with the annotated LAT values as local maxima of *dV* exceeding a threshold *dV*_th_. The third panel shows the resulting phases of the optical mapping signal.

For the simulations where the signals represent transmembrane voltages, these LATs are annotated using a voltage threshold *V*_th_ = − 60 mV [25]. The optical mapping signals represent unipolar voltages, but contain considerable signal drift. As such, LATs are annotated as the local maxima of the time derivative of the signal exceeding *dV*_th_ = 0 mV [28]. The clinical CARTO data contained pre-calculated LATs, so no conversion was needed. These LATs are then transformed into phases using sawtooth mapping [30], which linearly interpolates the phase between two consecutive LATs. Importantly, points activated fewer than two times are considered inactive and are assigned an undefined phase.

#### Phase index

The phase index is a topological property defined on a closed curve within a phase map. It quantifies the net change in phase around the curve and can be formally given by:

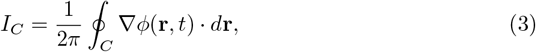

where *C* is a closed curve and *ϕ*(**r**, *t*) is the phase along *C*. In contrast to previous formulations that require the phase to be differentiable except at isolated points [5, 31], this work only requires the phase to be continuous and piecewise differentiable along the integration path. This allows the phase field to exhibit wavefront collisions, where the derivative of the phase is undefined, while still ensuring that the phase varies continuously along the integration path. Since the phase must return to the same state after a full traversal of the closed curve, the total change in phase around *C* must be an integer multiple of 2*π*. Consequently, the phase index *I*_*C*_ can only take integer values.

In electrophysiology, the phase index is used to identify certain activation patterns. Functional reentry, for instance, is typically associated with small curves that exhibit a non-zero phase index, indicating the presence of a phase singularity [4]. Similarly, anatomical reentry can be characterized by evaluating curves that encircle anatomical structures, and calculating whether their phase index is non-zero [32].

However, in cardiac tissue, the assumption of the phase map being continuous is not always valid. Regions may exist where *ϕ*(**r**, *t*) exhibits discontinuities, rendering the phase index ill-defined for curves that intersect them. Such regions are referred to as *phase defects* [21], which will be discussed in the following section.

#### Phase defects

Phase defects can be broadly categorized into three types: functional conduction block, structural conduction block, and boundaries, see Fig 5.

**Fig 5.**
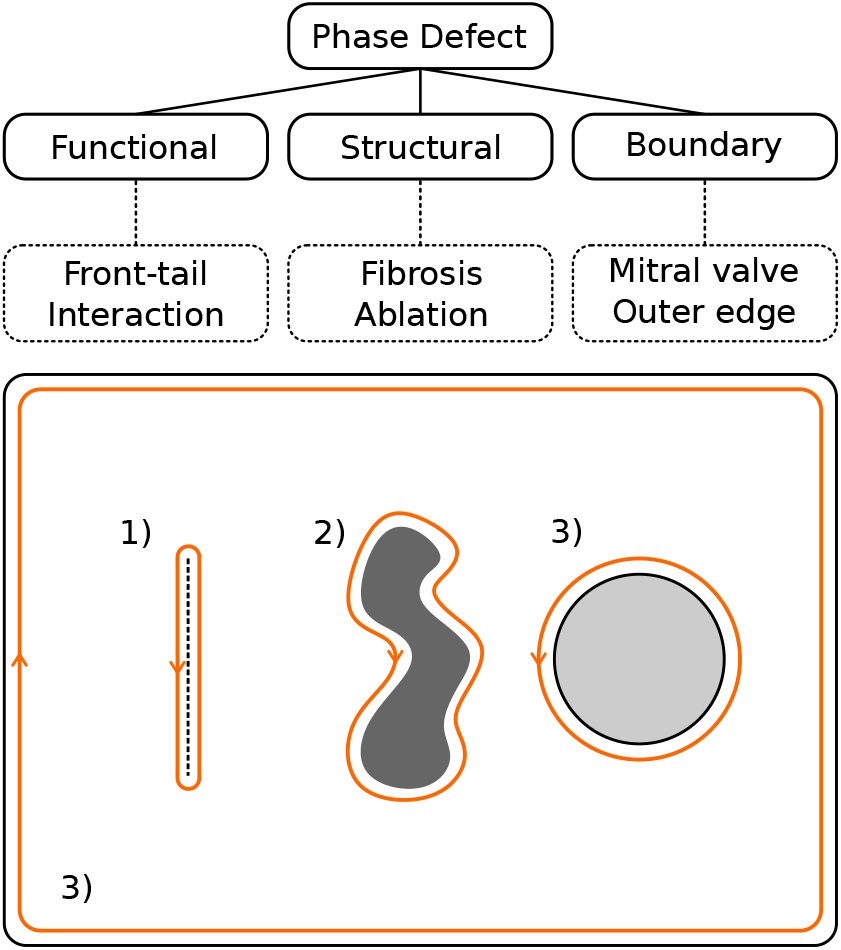
Illustration of functional, structural, and boundary phase defects. Illustration of the three types of phase defects on cardiac tissue: (1) Functional, (2) Structural, and (3) Boundary. An orange closed curve is shown around each phase defect, indicating the ideal shape of the closed curve C used to calculate the phase index. The orange arrow indicates the direction in which the phase index needs to be integrated, following established conventions [31]. Note that there are two boundaries present: a circular hole, and the outer edge of the rectangular tissue.

First, **functional conduction block** occurs when an excitation wavefront encounters tissue that is still in refractoriness. At such sites, one side of the interface is excited while the other is repolarizing, forming a line of discontinuity in the phase map [18, 21].

Second, **structural conduction block** can be caused by the non-conductive structures in the cardiac tissue, such as diffuse fibrosis or dense scar tissue that is created during ablation therapy. In transmembrane voltage maps obtained from simulations, structural conduction block manifests itself as undefined regions in the phase, when the cell voltage does not exceed the activation threshold. However, in electrogram data of optical mapping or electroanatomic mapping, the transmembrane voltage of small regions of cardiac cells gets averaged and smoothed. Therefore, when healthy tissue is surrounded by fibrotic tissue, the resulting electrogram will be affected. As such, structural conduction block would manifest itself as regions of discontinuous phase in electrogram data, akin to functional conduction block.

Third, **boundaries** refer to anatomical or geometric edges in the tissue, such as the borders of a 2D simulation or structures such as the MV, LPV or RPV [32].

To ensure that the phase index is well-defined, curves must avoid crossing discontinuous or undefined regions. Instead, the entire phase defect should be enclosed within the curve, as illustrated in Fig 5. The direction of integration follows established conventions [31], which are also indicated in the figure.

#### Conservation of total phase index

In case the phase map on the surface *S* contains a finite number of isolated singular points, there exists the following conservation law [31]:

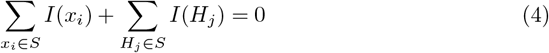

where *x*_*i*_ are the singular points in *S*, and *H*_*j*_ the holes in the surface *S*. This statement does not hold when conduction blocks are present within the phase map. However, it is possible to generalize Eq 4 by treating all phase defects as holes in a reduced surface *S*^′^ ⊂ *S*.

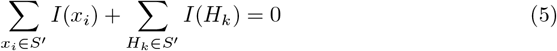

where *x*_*i*_ are the singular points in *S*^′^, and *H*_*k*_ the holes of *S*^′^, including all excised phase defects. From now on we will refer to Eq 5 as the index theorem.

#### Naive implementation of phase mapping

After constructing the phase map *ϕ*(**r**, *t*), the cardiac tissue is tessellated, and the phase index is computed for each mesh face at each time step. By naively assuming the phase map to be continuous, Eq 3 can be calculated by summing phase differences along the edges of each mesh face:

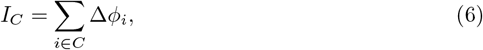

where Δ*ϕ*_*i*_ represents the circular difference between *ϕ*_*i*+1_ and *ϕ*_*i*_, i.e., the shortest path between two points on the unit circle. For most edges, this circular difference is simply Δ*ϕ*_*i*_ = *ϕ*_*i*+1_ − *ϕ*_*i*_. However, when the phase jumps from *π* to − *π* or vice versa, Δ*ϕ*_*i*_ must be adjusted by ± 2*π* to account for the periodicity of the phase. For a closed curve *C*, each phase value *ϕ*_*i*_ appears twice in the sum: once with a positive sign and once with a negative sign. As a result, the contributions of most edges cancel out, except for those where a phase jump occurs. Assuming the phase map is continuous, only edges with phase jumps will have an absolute phase difference larger than *π*. A positive jump is recorded when *ϕ*_*i*+1_ − *ϕ*_*i*_ *<* − *π*, and a negative jump when *ϕ*_*i*+1_ − *ϕ*_*i*_ *> π*. The phase index is then given by the difference in positive and negative jumps:

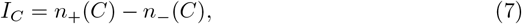

where *n*_+_(*C*) and *n*_−_ (*C*) are the number of positive and negative jumps along *C*, respectively.

Additionally, if a closed curve *C* passes through points with undefined phase, the phase index *I*_*C*_ cannot be meaningfully evaluated and is considered undefined.

In this implementation, phase singularities are the smallest closed contours with a non-zero phase index, representing loops that encircle points where the cardiac activity rotates.

#### Extended implementation of phase mapping

When phase defects are present in the cardiac tissue, the assumption of a continuous phase map is violated, and additional steps are required to ensure that Eq 3 is well-defined.

First, phase defects are identified and excluded from the mesh. To detect discontinuities, the absolute value of the circular phase difference | Δ*ϕ* | is evaluated across mesh edges [26]. The rationale is that | Δ*ϕ* | remains small where the phase map is continuous, but becomes large when an edge crosses a discontinuity. Faces containing at least one edge where | Δ*ϕ* | exceeds a predefined threshold are removed from the mesh, effectively creating holes. Thresholds of 0.1*π*, 0.4*π*, and 0.2*π* were used for simulated, experimental, and clinical data, respectively. These values were selected to ensure accurate annotation of regions where conduction block was expected. Nevertheless, low threshold values should be used with caution, as they can lead to slow-conducting regions being misclassified as conduction blocks due to the high local phase gradient. In addition, mesh faces containing at least one node with an undefined phase are also removed.

Second, the boundaries of the remaining mesh are extracted to form the closed contours *C* of various sizes, which are then used to evaluate the phase index using Eq 7.

Finally, the remaining mesh is evaluated according to the procedure described in Section, providing the smallest possible closed contours equal to the mesh faces.

In this implementation, we define critical phase defects as the phase defects with a non-zero phase index, which represent regions of rotational activity [20, 32].

## Results

### Finitewave simulations

To illustrate the key features of each simulation, results are shown for a representative single time frame. The full analysis code and videos are provided as supplementary material (see Section), enabling inspection of other time frames.

#### Linear rotor core

Fig 6 shows a simulation of functional reentry, with a core that displays distinct rotational activity around a phase defect. The absence of non-conducting regions in the action potential field suggests that the defect represents a functional conduction block. The naive approach identifies one or multiple phase singularities of both positive and negative phase index values at the central defect per time frame. In contrast, the extended approach finds two critical phase defects per time frame: the central defect a phase index of minus one, and the outer boundary with a phase index of one. Moreover, the extended approach adheres to the index theorem for all time frames in contrast to zero time frames for the naive approach. This result aligns with the underlying tissue dynamics: namely, a single central rotation, which is also clearly visible in the supplementary video (S1 Video).

**Fig 6.**
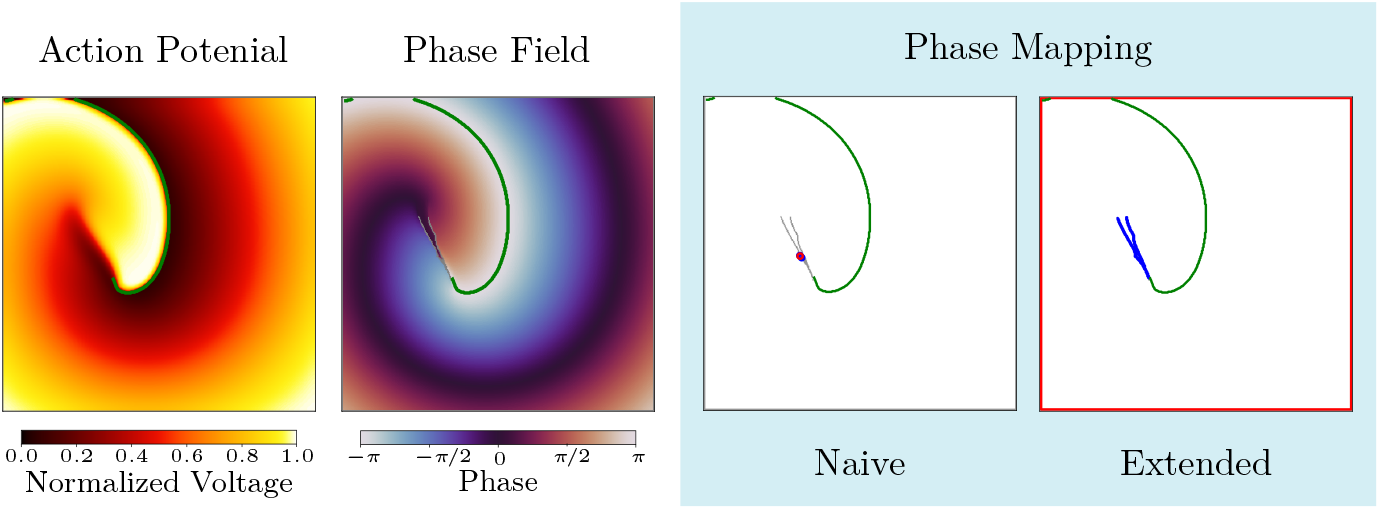
Phase mapping analysis of a linear core simulation. Snapshot (t=594 ms) of the analysis of the simulation of a linear core using the Fenton-Karma model. The wave front is annotated in green, and the phase defects are annotated in gray. *First panel*: Normalized action potential. *Second panel*: The phase map constructed from the action potential. *Third panel*: The results of the naive phase mapping. The blue and red circles denote phase singularities with phase indices of minus and one, respectively. *Fourth panel*: The results of extended phase mapping. The blue and red lines denote closed curves with phase indices of minus one and one, respectively.

#### Parallel activation around a functional conduction block

Fig 7 illustrates a functional conduction block in the absence of rotational activity. As in the previous case, the action potential field shows no non-conducting regions, indicating that the phase defect is again a functional conduction block. Despite the lack of rotation, the naive approach detects an average of four phase singularities around the defect per time frame. In contrast, the extended approach finds an average of zero critical phase defects and seven non-critical phase defects per time frame. Additionally, the phase index is conserved for all time frames in the extended approach, in contrast to 96.62% for the naive approach. Given that the simulation exhibits no rotations, as evident from both the snapshot and the supplementary video (S1 Video), extended phase mapping is the only approach that produces results consistent with the non-rotational nature of the simulation.

**Fig 7.**
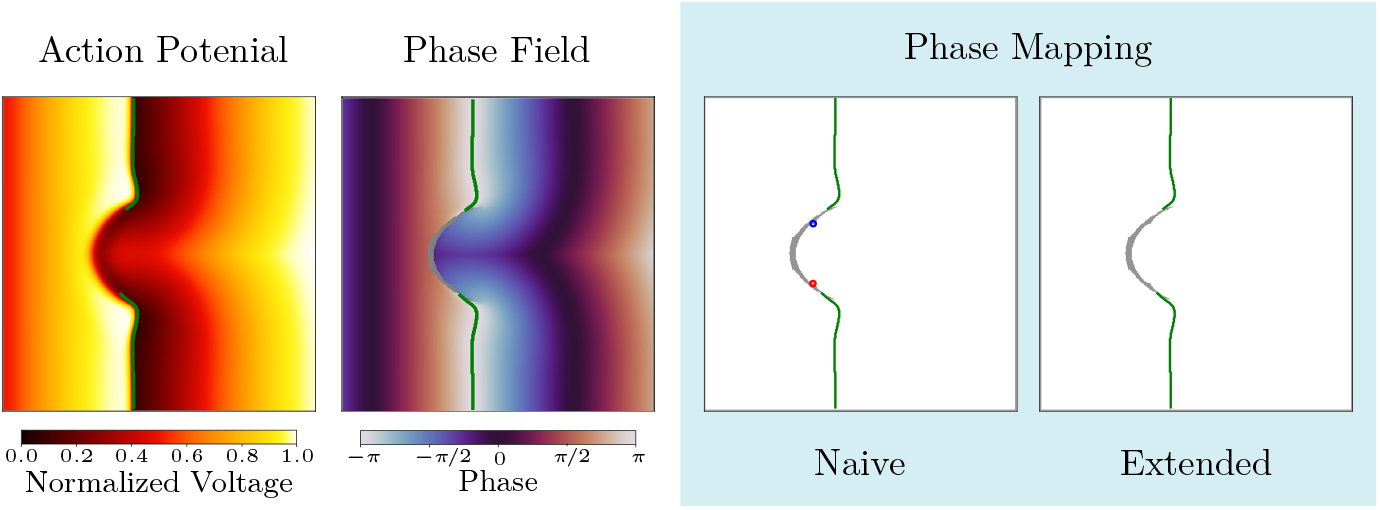
Phase mapping analysis of a functional conduction block simulation. Snapshot (t=722 ms) of the analysis of the simulation of a functional conduction block using the Fenton-Karma model. The wave front is annotated in green, and the phase defects are annotated in gray. *First panel*: Normalized action potential. *Second panel*: The phase map constructed from the action potential. *Third panel*: The results of the naive phase mapping. The blue and red circles denote phase singularities with phase indices of minus and one, respectively. *Fourth panel*: The results of extended phase mapping. The blue and red lines denote closed curves with phase indices of minus one and one, respectively.

#### Complete and near-complete rotation around scar tissue

Finally, Fig 8 shows two simultaneous rotations around obstacles of different sizes. Unlike the previous simulation, the phase defects in this case are structural conduction blocks, which are visible in both the action potential field and the phase map. The snapshot and supplementary video (S1 Video) demonstrate that the smaller obstacle supports a complete rotation, while the larger scar shows a near-complete rotation that is interrupted before the wavefront can fully circle the obstacle. The naive approach detects zero phase singularities per time frame, since the phase within scar tissue is undefined. The extended approach identifies an average of two critical phase defects per time frame: the smaller one receives a phase index of one, and the larger one a phase index of minus one. Additionally, the tissue boundary is identified as a non-critical phase defect for all time frames. Furthermore, the naive and extended approaches adhere to the index theorem for all time frames.

**Fig 8.**
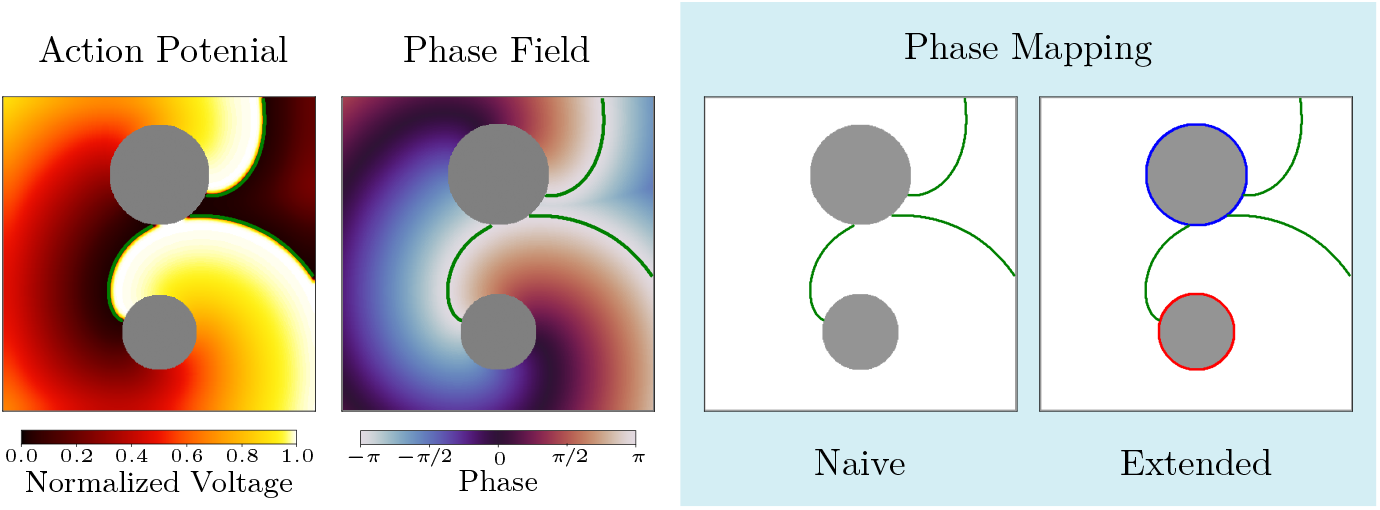
Phase mapping analysis of a reentry pair with complete and near-complete rotation. Snapshot (t=906 ms) of the analysis of the simulation showing a reentry pair with complete and near-complete rotation using the Fenton-Karma model. The wave front is annotated in green, and the phase defects are annotated in gray. *First panel*: Normalized action potential. *Second panel*: The phase map constructed from the action potential. *Third panel*: The results of the naive phase mapping. The blue and red circles denote phase singularities with phase indices of minus and one, respectively. *Fourth panel*: The results of extended phase mapping. The blue and red lines denote closed curves with phase indices of minus one and one, respectively.

### Optical mapping of ventricular fibrillation

The differences between the naive and extended phase mapping approaches become even more pronounced when applied to optical mapping data of ventricular fibrillation, illustrated in Fig 9. This case is characterized by multiple activation wavefronts interacting with each other, hosting both stable and unstable rotational centers, characteristic of fibrillation. The naive approach produced an average of 37 static and meandering phase singularities per time frame, obscuring the underlying activation pattern. In contrast, the extended approach clarified the analysis by detecting an average of 12 critical phase defects and 17 non-critical phase defects per time frame. Furthermore, the naive approach adhered the index theorem for 18.66% of the recorded frames in contrast to 99.97% for the extended approach. Additionally, the extended approach discovered the presence of critical phase defects hosting more than one wavefront. Moreover, the phase defect containing the outer edge occasionally had an absolute phase index larger than one due to the presence of multiple wavefronts. This indicates that more complex activation patterns are present than previously described in the simulations, where the phase index was always equal to minus one, zero, or one.

**Fig 9.**
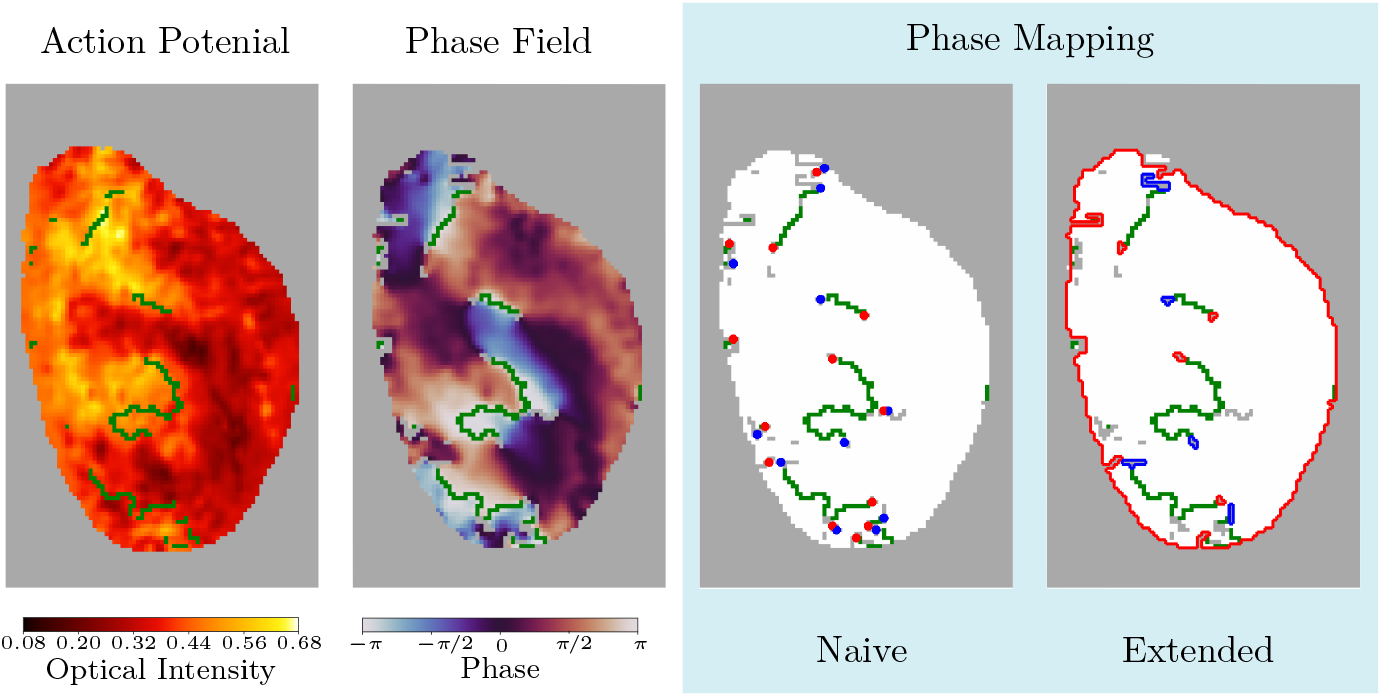
Phase mapping analysis of optical recordings during ventricular fibrillation. Snapshot (t=293 ms) of the analysis of an experimental optical mapping recording of ventricular fibrillation in a rat heart. The wave front is annotated in green, and the phase defects are annotated in gray. *First panel*: Optical intensity map. *Second panel*: The phase map constructed from the optical map. *Third panel*: The results of the naive phase mapping. The blue and red circles denote phase singularities with phase indices of minus and one, respectively. *Fourth panel*: The results of extended phase mapping. The blue and red lines denote closed curves with positive and negative phase indices, respectively.

### CARTO map of clinical atrial tachycardia

The clinical case in Fig 10 clearly illustrates several key differences between the naive and extended phase mapping approaches. This case was clinically diagnosed as atrial tachycardia sustained by reentry around the MV + LPV and a nearby scar line. The tachycardia was terminated via ablation between the MV and the scar. The naïve approach detects 21 static phase singularities in the proximity of the phase defects as detected by the extended method, which is poorly interpretable and far from realistic expectations. The anatomical valves, including the MV and LPV, are also completely devoid of any detections, suggesting that the naive approach does not result in the correct diagnosis. In contrast, the extended approach shows a different result. Two static critical phase defects were consistently detected around the MV + LPV and a nearby phase defect line. Additionally, 35 static non-critical phase defects are consistently annotated, indicating the presence of conduction block. In particular, the previous ablation line between the MV and LPV was correctly annotated as a conduction block. These results are in line with the diagnosis, where the MV + LPV and an anterior scar were found to host reentry. Additionally, this agrees with the index theorem previously described in [20, 32], stating that the sum of all phase indices must equal zero. This is in contrast to the naive approach, which violates the index theorem for every time frame of the recording.

**Fig 10.**
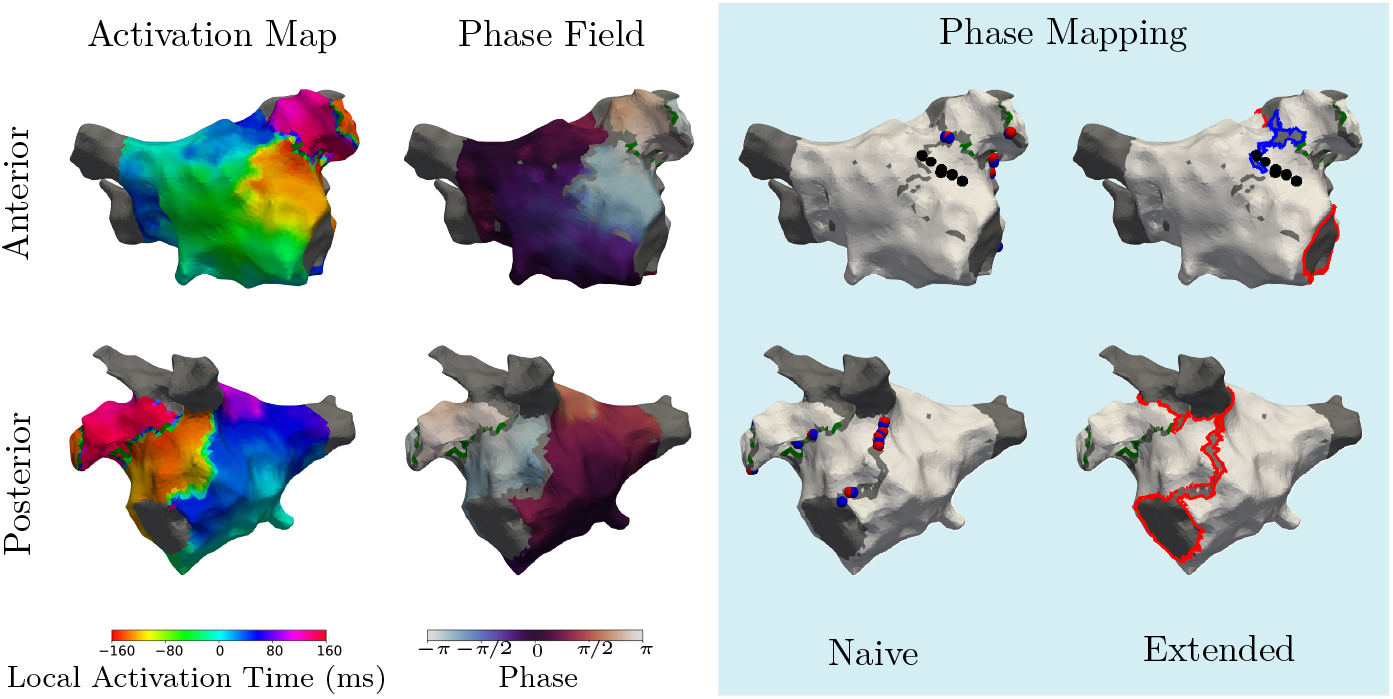
Phase mapping analysis of a clinical atrial tachycardia. Snapshot (t=0 ms) of the clinical CARTO map of atrial tachycardia diagnosed with reentry around the MV + LPV and a nearby scar. The wave front is annotated in green, the phase defects are annotated in gray, and the ablation points are annotated in black. *First panel*: Local activation time map. *Second panel*: The phase map constructed from the LAT map. *Third panel*: The results of the naive phase mapping. The blue and red spheres denote phase singularities with phase indices of minus and one, respectively. *Fourth panel*: The results of extended phase mapping. The blue and red lines denote closed curves with phase indices of minus one and one, respectively.

## Discussion

### Main findings

This section interprets the results presented above, highlighting the shortcomings of naive phase mapping across simulated, experimental, and clinical data, and how these are mitigated by the extended approach.

#### Shortcomings of the naive phase mapping approach

The first simulation, featuring a linear rotor core (Fig 6), shows that the naive approach detects both static and meandering phase singularities around the central phase defect. One could argue that clustering these singularities might still provide a correct interpretation. Indeed, post-processing techniques such as clustering have been shown to improve phase mapping in several studies [24, 33, 34]. However, the extended approach highlights the shape of the reentry core, offering a more elegant solution.

Moreover, in the second simulation showing parallel activation around a phase defect, clustering the naive phase mapping results (Fig 7) would still incorrectly identify the phase defect as two opposite phase singularities. To mitigate such false detections, one could require that only detections associated with full rotations should be considered. This can be done by requiring a minimal duration that singular points should exist, effectively filtering out transient or non-rotational patterns [33]. However, this again requires additional processing, which is unnecessary in the extended approach.

Finally, the third simulation, involving complete and near-complete rotation around scar tissue (Fig 8), challenges the phase singularity filtering strategy for naive phase mapping based on the number of rotations or duration. The naive approach detects no phase singularities due to the undefined phase on the scar tissue, indicating an absence of rotation. Other implementations of phase mapping detect all wavefront endpoints as phase singularities, irrespective of the nature of their anchor point [6]. However, for these implementations, the interpretation would still imply that the phase singularities surrounding the scar structures do not make a full rotation, meaning they would be disregarded by the previously mentioned phase singularity filtering strategy. In these cases, the extended approach invariably produces superior results

In the optical mapping data, the presence of most phase singularities shows a similarity to the second simulation of parallel activation around a phase defect, where many false positive phase singularities were also identified. This indicates that the false positive detections of the naive approach were largely due to parallel activation around a phase defect, akin to the second simulation depicted in Fig 7. Another explanation for the presence of phase defects is signal noise or low mesh resolution, as this can manifest itself as discontinuous regions in the phase map.

The clinical case similarly demonstrates the poor interpretation of the naive approach in a realistic setting. The naive approach finds many false positive detections due to the presence of many phase defects, ranging from anatomical valves to possible scar tissue or functional block.

In conclusion, the naive method ignores large non-conductive obstacles, and all detected singular points are located around phase defects. This last observation strongly indicates that the phase singularities arise from the presence of phase defects, rather than rotational activity around singular points. For each individual simulation, the naive approach could still be interpreted reasonably when combined with the necessary pre- and postprocessing tailored to the data. However, when interpreting all five case studies together, it is clear that naive phase mapping does not provide a consistent interpretation that is in accordance with physiological expectations. The extended approach directly addresses these issues by redefining how phase discontinuities and non-conductive regions are treated. The following section illustrates how this resolves the inconsistencies observed with the naive method.

#### Proper handling of phase defects improves the detection of critical rotations

For the first simulation, the extended approach detects a single dynamic central critical phase defect denoting the linear rotor core and a single static critical phase defect denoting the outer edge of the tissue. This is interpreted as paired reentry, with a central semi-stable complete rotation and a stable near-complete rotation at the boundary.

The second simulation reveals a dynamic central non-critical phase defect, in addition to a static non-critical phase defect consisting of the outer edge of the tissue. The interpretation is that the excitation wave hits a functional conduction block, breaks, and finally recombines, meaning no rotational activity is involved.

In the third simulation, two critical phase defects and a non-critical phase defect are found, which are interpreted as a near-complete rotation and a complete rotation around the central scar structures, and no rotation around the outer edge. This is similar to the first simulation and differs only in the nature of the phase defects. Although the boundary in the first simulation and the larger scar structure in the third simulation do not support a full rotation, the extended approach still assigns them a nonzero phase index. Initially, such patterns were considered clinically irrelevant due to the absence of physical rotation, and were occasionally referred to as ‘wannabe-reentries’ [35]. However, recent clinical studies have shown that near-complete rotation can play a critical role in guiding ablation strategies [20, 36, 37]. This finding underscores the importance of recognizing activation patterns beyond full rotation, and highlights the limitations of relying solely on phase singularity clustering and/or filtering strategies.

The optical mapping data results give the impression that there are only several rotational drivers sustaining the fibrillation, characterized by one or multiple activation wavefronts, agreeing with previous results described in [16]. These rotational drivers are described by phase defects with a non-zero index, which can be stable or unstable.

Lastly, the clinical data analysis interprets this activation map as paired reentry around the MV and LPV, as well as around an anterior scar. This finding is consistent with the clinical diagnosis. Additionally, the identification of a pair of counter-rotating waves results in a total phase index of zero, thereby adhering to the index theorem [31], further supporting the consistency of the extended approach in identifying rotational drivers across diverse datasets.

In conclusion, the extended approach provides a physiologically consistent interpretation across all cases by explicitly accounting for phase defects. Rotational drivers are identified as static or dynamic phase defects, each characterized by a non-zero phase index. In contrast, the naive approach fails to detect these rotations, in all given examples.

### Previous research

This paper is closely tied to previous research concerning the detection of phase defects and rotational drivers.

Several previous studies have addressed the role of phase defects in the analysis of activation maps. Notably, [18, 21, 26] emphasized the importance of phase defects when interpreting phase maps.

In particular, [26] pointed out that the phase index is ill-defined in the proximity of phase defects, implying a limitation of naive phase mapping. The results in Fig 7 indicate the importance of assigning a phase index to phase defects to distinguish between parallel activation and rotational activity.

Furthermore, [18] argues that the center of a spiral wave is more appropriately interpreted as a phase defect with a finite circumference rather than as a singular point. Likewise, our results support this interpretation: once the interactions between the wavefront and the refractory tail are properly accounted for by detecting phase defects, the apparent singular points vanish, as illustrated in Fig 6 and Fig 7.

The influence of phase defects has been indirectly addressed in previous studies, such as [24] and [34], which identified phase jumps by setting the phase-difference filter threshold above *π*. This approach effectively filters out the fraction of conduction blocks with an absolute phase difference above *π*, but it may still handle large non-conducting regions and boundaries improperly. Moreover, detecting phase defects and phase jumps in a single step, restricts the use of additional features to improve phase defect detection.

In related work, several methods have been proposed to characterize the nature of non-excitable structures. [36, 37] reported observations of critical anatomical features exhibiting patterns similar to those shown in Fig 8. However, more notably, [20, 32] employed the same phase index used in this manuscript to assess the criticality of anatomical boundaries (e.g., MV, LPV, RPV). Building on this research, we extend the approach by incorporating all kinds of phase defects.

### Limitations and future prospects

The primary aim of this work is to highlight the challenges that arise when phase defects are present and to demonstrate the potential benefits of explicitly addressing them during phase mapping analysis. As such, this study should not be viewed as a detailed statistical analysis on a large dataset investigating phase defect detection algorithms and robustness. We will now describe four limitations that are tied to this.

First, the method used to detect phase defects in this study, relying on one parameter, is relatively basic and intended primarily for demonstration purposes. To make phase mapping more reliable and robust for interpreting realistic data, more sophisticated approaches relying on more features of the electrogram for phase defect detection should be explored. For example, features such as low voltage, mid-diastolic potentials, after-depolarizations, double potentials, fractionation, and other electrogram characteristics associated with decreased cardiac conduction [38, 39] could provide more robust indicators of phase defects and are valuable future research topics.

Second, the size of our dataset is severely limited to five cases. The simulations used in this work are also intentionally simplified and do not reflect realistic cardiac tissue. The reason for this was to isolate the core problems with the naive implementation of phase mapping, removing additional effects such as high signal noise and low mesh resolution. This limitation is partly mitigated by including the analysis of the optical map and clinical activation map, reflecting more realistic scenarios. Our research group is currently working on analyzing larger datasets of atrial and ventricular tachycardia using extended phase mapping to obtain more statistically significant and robust results.

Third, the effects of signal noise and mesh resolution were not investigated in this study. The effect of these factors on phase mapping has been demonstrated in previous studies [7, 8, 14]. In this work, all simulations contained a very high resolution and no signal noise; therefore, we excluded the effects of these factors on the interpretations of these results. This could not be done for the experimental data, as the optical map contained a relatively low resolution and considerable signal noise [27], possibly impacting our results. Similarly, the clinical activation map had a high resolution, but contained interpolation artifacts that manifested themselves as noise in the resulting phase map. Consequently, we acknowledge the effect of signal noise and resolution on our results, and propose their investigation as a valuable future research topic.

Finally, the influence of the definition of phase on our results was not investigated. In this paper, we refrained from using the conventional phase definition based on the

Hilbert Transform due to erroneous phase defect detections near wavefronts [26], leading to incorrect interpretations. As such, we opted for a modified definition of phase recommended by [26] based on an activation threshold and consecutive activation times, yielding a sawtooth-shaped phase. We acknowledge the influence of this activation threshold on the interpretation of our results, and leave the investigation of its influence for future studies.

## Conclusion

We demonstrated the influence of phase defects on the detection of phase singularities using a naive phase mapping implementation across five case studies encompassing simulated, experimental, and clinical data. Comparing this naive implementation to an extended approach that explicitly detects critical phase defects, we found that the naive method yields inconsistent interpretations across cases. In contrast, the extended approach provides consistent results and interprets the drivers of rotation as phase defects characterized by a non-zero phase index. These findings challenge the classical view that rotors or functional reentry in cardiac arrhythmia are sustained by physical rotation around singular points. Instead, we propose that arrhythmia drivers arise from complete or near-complete rotation around phase defects. Rather than promoting our specific implementation, we emphasize the necessity of explicitly addressing phase defects in phase mapping analyses. We hope this work motivates further research into the role of phase defects and encourages their integration into analytical workflows to improve understanding of complex activation patterns.

## Supporting information

**S1 Video. Supporting videos**. All supplementary videos can be found on Zenodo (http://doi.org/10.5281/zenodo.17602155).

**S1 Data. Optical Mapping Data**. The optical data that is presented and analysized in this paper can be found on Zenodo (http://doi.org/10.5281/zenodo.18319101).

**S1 Code. Code Repository**. Readers who wish to reproduce our results can find the code and usage instructions in our GitLab repository (https://gitlab.com/opendgm/paper-phase-defects).

## Acknowledgments

We thank Balvinder Handa, Vineesh Kappadan, and Fu Siong Ng for providing optical mapping data, and Mattias Duytschaever for providing the clinical case.

